# High-throughput Analysis of Arabidopsis Stem Vibrations to Identify Mutants with Altered Mechanical Properties

**DOI:** 10.1101/315838

**Authors:** Miyuki T. Nakata, Masahiro Takahara, Shingo Sakamoto, Kouki Yoshida, Nobutaka Mitsuda

## Abstract

Mechanical properties are rarely used as quantitative indices for the large-scale mutant screening of plants, even in the model plant *Arabidopsis thaliana*. The mechanical properties of plant stems generally influence their vibrational characteristics. Here, we developed Python-based software, named AraVib, for the high-throughput analysis of free vibrations of plant stems, focusing specifically on Arabidopsis stem vibrations, and its extended version, named AraVibS, to identify mutants with altered mechanical properties. These programs can be used without knowledge of Python and require only an inexpensive handmade setting stand and an iPhone/iPad with a high-speed shooting function for data acquisition. Using our system, we identified an *nst1 nst3* double-mutant lacking secondary cell walls in fiber cells and a *wrky12* mutant displaying ectopic formation of secondary cell wall compared with wild type by employing only two growth traits (stem height and fresh weight) in addition to videos of stem vibrations. Furthermore, we calculated the logarithmic decrement, the damping ratio, the natural frequency and the stiffness based on the spring-mass-damper model from the video data using AraVib. The stiffness was estimated to be drastically decreased in *nst1 nst3*, which agreed with previous tensile test results. However, in *wrky12*, the stiffness was significantly increased. These results demonstrate the effectiveness of our new system. Because our method can be applied in a high-throughput manner, it can be used to screen for mutants with altered mechanical properties.

## Introduction

Lignocellulosic biomass, which comes mainly from plant cell walls, is one of the largest renewable resources, which are indispensable for human society. It has an extremely broad range of uses, including in fuels, paper, building materials, furniture, crafts, biochemicals, biomaterials and acoustic instruments. Consequently, its production and the selection of plant species with high-yield/quality lignocellulose levels are being actively studied. Lignocellulosic biomass consists mainly of cellulose, hemicellulose and lignin, which are high-molecular weight polymers that interact with each other in a complex manner, making lignocellulose highly recalcitrant. While primary cell walls exist in all plant cells, thickened secondary cell walls are preferentially formed in vessel and fiber cells to confer hydrophobicity and robustness to tissues, contribute to the transport of water and nutrients, support plant bodies and prevent pathogen invasion. Because the secondary cell walls form a large proportion of the plant biomass, their recalcitrance to degradation could be a focus of molecular and traditional breeding.

In model plants, the large-scale screening of various mutants and transformed strains in which lignocellulose could be modified has been attempted. In particular, *Arabidopsis thaliana* is often used as the model plant because it has many advantages, such as a short-generation period, and the feasibility of genetic engineering and forward genetic screening. In a phenotypic screening, anatomical changes, biomass and the cellulase saccharification efficiency have been mainly used as indices (Santoro et al., 2010; Wang et al., 2010; Cassan-Wang, 2013; Sakamoto and Mitsuda, 2015). However, even though the mechanical properties of the stem are important in physiological studies and applications, their evaluation in a large-scale screening usually relies on primitive and qualitative methods, such as visual observations and hand-bending tests. However, other mechanical tests, such as tensile, compression and mechanical bending tests (Gibson, 2012), can be applied to *A. thaliana* (Ryden et al., 2003; MacMillan et al., 2013; Sakamoto et al., 2016; Brulé et al., 2016). Micro-mechanical or micro-tensile tests combining deformations and monitoring have also been established (Mott et al., 1996; Köhler and Spatz, 2002). The methods for investigating the strength of stem in cellular and/or cell-wall level, as well as tissue level, are available. Additionally, in recent years, a method measuring local mechanical properties using an atomic force microscope has been developed (Peaucelle et al., 2011; Radotić et al., 2012; Fernandes, 2012; Braybrook and Peaucelle, 2013; Peaucelle, 2014). However, these assays are not commonly used for screening because they require expensive and specialized stationary instruments. To find useful genes and mutations with high industrial values, the mechanical properties of the stems could be important indices. Thus, the development of a high-throughput and low-cost quantitative screening method is required.

In this study, we focused on the stem’s vibrational frequency as an indicator of its mechanical properties. Vibration is a general physical phenomenon observed when an object is deformed by an external force. Every object has its own vibrational frequency, which depends on its own stiffness and mass. In fact, vibrational characteristics are used for evaluating mechanical properties in various plants, and various methods for measuring vibration have been proposed. For instance in wood, hitting it with a hammer and analyzing sound waves is generally used for calculating the mechanical properties (Yamamoto et al., 1980; Halabe et al., 1997). Additionally, the resonance method also has been used to examine the mechanical properties of plant tissues (Finney and Abbott, 1979; Virgin, 1955; Falk et al., 1958; Burström et al., 1967; Uhrstörm, 1969; Burström et al., 1970; Burström, 1971; Uhrstörm and Svensson, 1979; Cosgrove and Green, 1981; Niklas and Moon, 1988). The flexural rigidity of the stem has been evaluated using free vibrations (Żebrowski, 1991; Spatz and Speck, 2001). Even in *A. thaliana*, a modal analysis of stem vibration was attempted (Der Loughian et al., 2014), and the possibility of a nondestructive phenotyping method was proposed (Shah et al., 2017). However, these studies aimed to calculate physical property values but did not intend to apply them to the large-scale screening of mutants with altered mechanical properties.

In this research, to promote the application of vibrational analyses in the field of molecular genetics using *A. thaliana*, we developed Python-based software for the high-throughput analysis of <u>Ara</u>bidopsis stem <u>vib</u>rations (named AraVib) and an extended version, named AraVibS, to identify Arabidopsis mutants with altered mechanical properties. The software are free of charge and can be used without any knowledge of Python. For data acquisition, we have established a method to analyze the free vibration of the inflorescence stems of *A. thaliana*. Here, we demonstrate that our system can rapidly estimate the mechanical properties of stems and discriminate mutants having altered cell walls.

## Materials and methods

### Plant strains and growth conditions

*A. thaliana* accession Columbia-0 was used as the wild type. The *nst1-1 nst3-1* double mutant, which has defects in secondary cell wall formation in fiber cells of the inflorescence stem (Mitsuda et al., 2007; Zhong et al., 2007), was described in Mitsuda et al. (2007). The *wrky12-1* mutant, in which the deposition of secondary cell wall components were ectopically observed in the pith cells, was described in Wang et al. (2010). The seeds were rinsed in a Plant Preservative Mixture solution (3% Plant Preservative Mixture and 0.1% Triton-X 100; Nacalai Tesque Inc., Kyoto, Japan), sown on a culture soil (soil mix:vermiculite, 1:1) and cultivated for ~1.5–2 months under long-day conditions (16-h light/8-h dark) at 22°C. In experimental condition 1, multiple plants were grown in the same plant pot, and the area per plant was 1,200 mm^2^ or less. In experimental condition 2, plants were grown individually in ARASYSTEM (http://www.arasystem.com/), and the area per plant was ~2,000 mm^2^.

### Imaging of stem vibrations

Our method consists of three sections: measuring growth traits, settings and imaging (Figures 1, 2) after harvesting the stem. First, we measured growth traits, including the stem height (H) and 10-cm stem fresh weight (FW). H was measured with a ruler. To reduce the effect of non-uniformity in weight distribution, the basal part of the stem was trimmed to a length of 10 cm with scissors, and all accessories, such as leaves, branches, flowers and fruits, were removed. The FW was weighed with a precision balance. Second, we set the 10-cm stem sample. The 1-cm position at the base was marked with a black marker, and the apical end of the stem was marked with red marker. To ensure detection by AraVib, a large enough amount of the red marker should be applied. The weight of the applied red marker itself was 1% or less of tested stems and, therefore ignorable. The 1-cm stem section below the black mark was sandwiched between plastic boards and firmly fixed with a rubber band and then placed on a plastic box (termed “a handmade setting board”). The plastic boards and the plastic box were made from refill attachments for 250-μL pipette tips (RAININ: GPS-250S). The background was covered with a black cloth or cardboard. The iPad/iPhone camera was fixed at a position 20 cm away from the stem. Third, we imaged stem vibrations. To apply linear bending, the upper position of the stem (3–4 cm from the tip) was grasped by tweezers and pulled sideways at an angle without breaking the stem. Immediately after the shooting had started, the stem was released (Supplementary Movies 1, 2). The image of the stem vibration was captured for 3–4 s with the high-speed shooting function (240 fps) of iPad/iPhone video. These procedures were repeated for each sample. Because the time from the measurement of the stem length to the end of capturing the vibration is within 2–3 min per sample (Figure 2), the change in FW resulting from a loss of moisture during the experimental process should be small.

**Figure 1.**
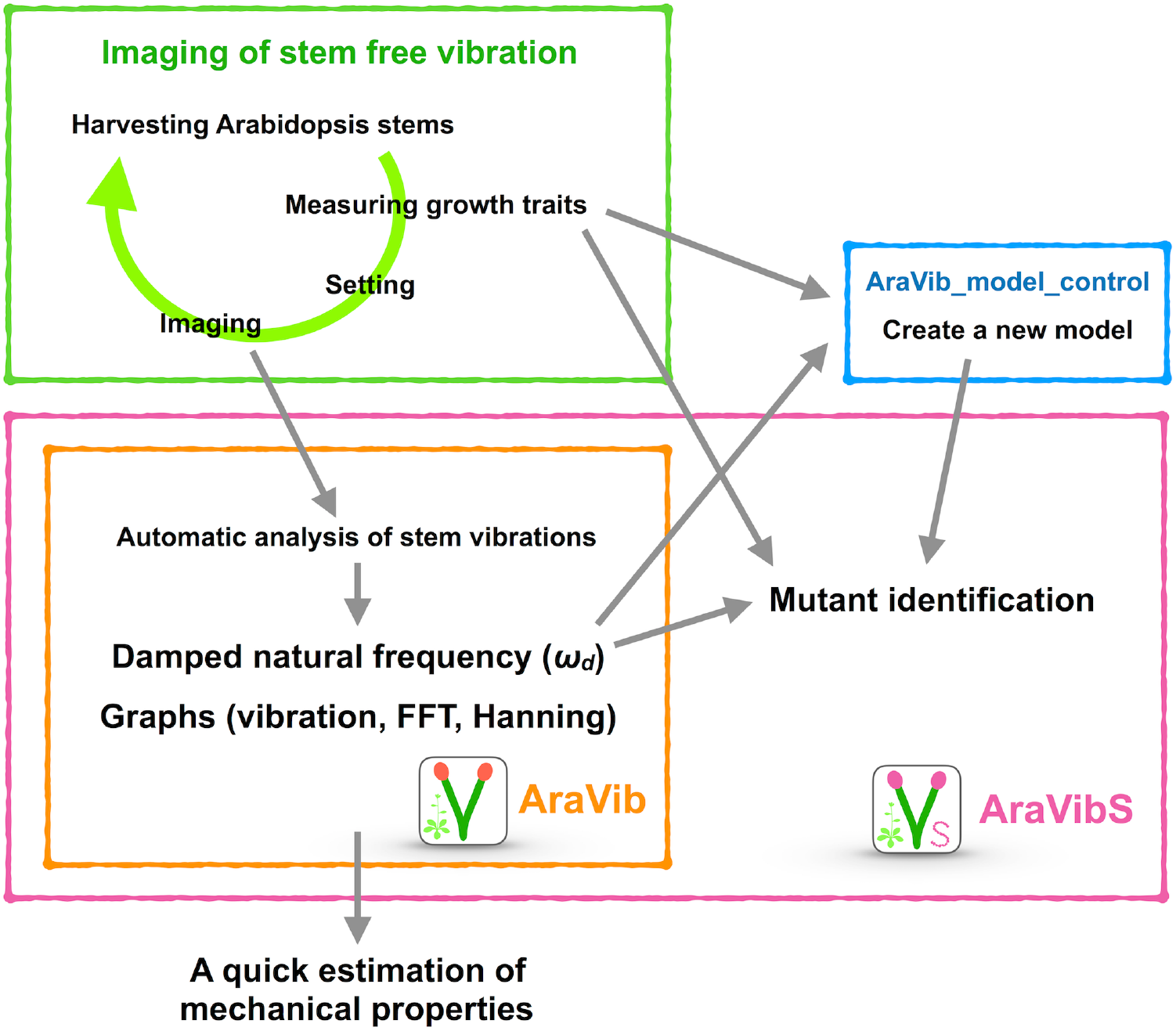
Schematic view of the high-throughput analysis of stem vibrations in this study. This analysis consists of the method for imaging of stem free vibration (green) and the AraVib software for automatic analysis of vibration (orange) and its extended version AraVibS for mutant identification (magenta). The AraVib_model_control program for automatically learning a model used for mutant identification (blue) is also included.

**Figure 2.**
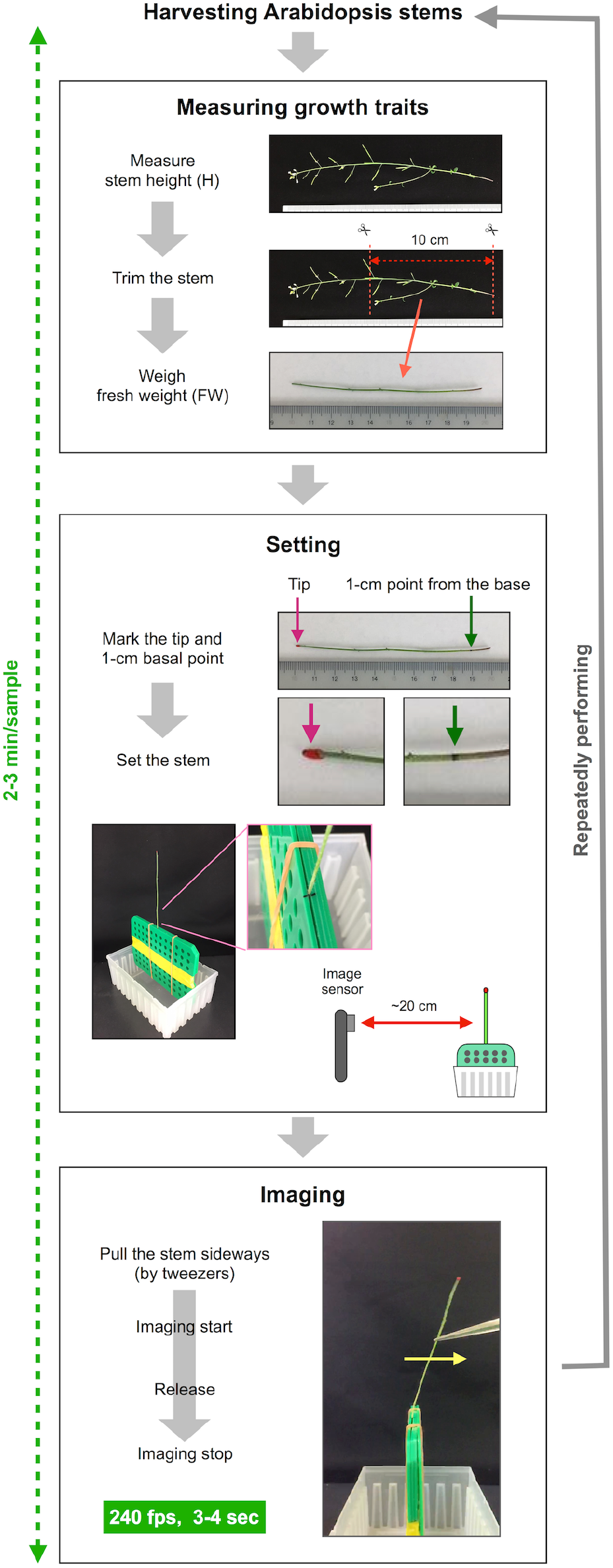
The procedure of imaging of stem free vibration. This method can roughly be divided into three steps besides harvesting stems. The first step is to measure growth traits. In this step, the stem height is measured, the basal part of the stem is trimmed to a length of 10 cm, the fresh weight of 10-cm stem is weighed. The second step is setting the stem sample. In this step, the trimmed stem is marked and set to the setting board. The setting board with the stem is placed ~20 cm from the image sensor of the iPhone/iPad. The third step is imaging. The stems are pulled sideways by tweezers, retained and released after imaging has started. Stem vibration is imaged in the slow-motion mode at 240 fps for 3-4 seconds. This series of procedures is repeated for each sample.

### Development of AraVib and AraVibS

AraVib is a command-line software that automatically analyzes the stem vibration movies, which were recorded using the above-mentioned method, in a batch (Figures 1, 3). In AraVib, the red marker on the black background is tracked, the centroid coordinates of the marker are determined, and the coordinates are converted to displacement data. The displacement data are then multiplied by the Hanning window (Harris, 1978), and the Hanning-multiplied data are used to calculate the damped natural frequency (*ω_d_*), using Fast Fourier Transform (FFT). Details of the algorithms and their optimization procedures for AraVib can be found in the Supplementary materials and Supplementary Figures 1–3. In this study, AraVib was run on the MacBook Pro (2.4 GHz Intel Core i7, 8GB 1600 MHz DDR3, macOS Sierra).

**Figure 3.**
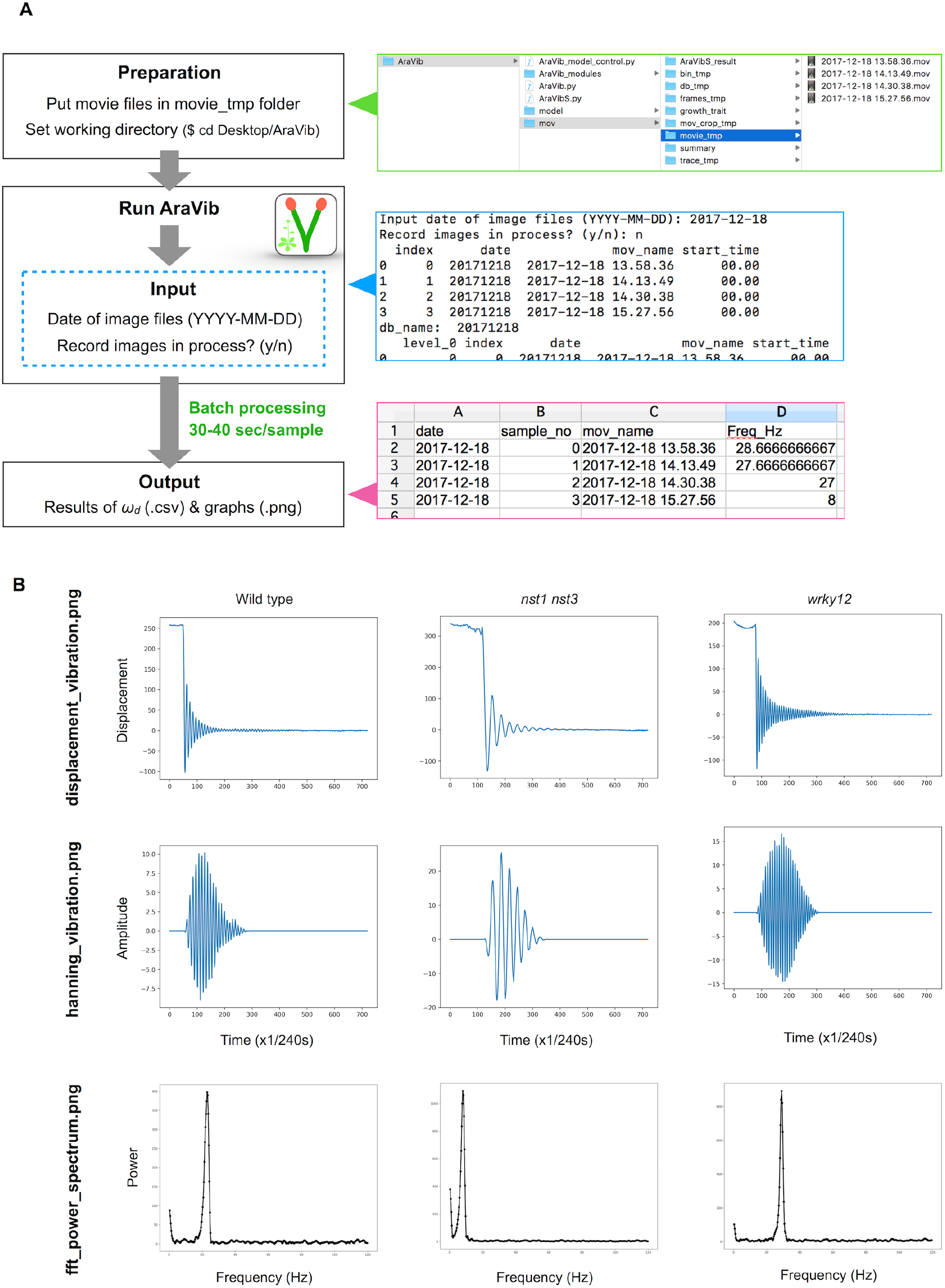
Outline of vibration analysis using AraVib. (A) Analysis diagram of the AraVib software. The procedure consists of three steps: preparation, run and output. For details, please refer to the GitHub page of AraVib. AraVib is capable of batch processing of all at once for each shooting date. In the upper right pannel is a folder structure, the right middle pannel shows an example of the command-line display, and the lower right pannel shows the output of *ωd* result. (B) Examples of three types of graphs to be output. displacement_vibration.png represents the graph of the raw vibration waveform. hanning_vibration.png represents a graph of the vibration waveform mutiplied by Hanning window. fft_power_spectrum.png represents a graph of the FFT power spectrum. See also Supplementary figure 4.

The latest version of the compressed AraVib folder containing these programs and some sample files is available in the GitHub page of AraVib. Details on how to use these programs are also available. The free Python packages or libraries, which are used in AraVib and AraVibS, are the OpenCV-Python package, NumPy package, the Python Pandas package, SciPy library, the pyplot package of the Matplotlib and Scikit-learn package. Details on how to set up them are shown in Supplementary materials.

The analytical flow of AraVib is shown in Figure 3A. The output is the csv file of the *ω_d_* list (Figure 3A) and three graphs of each samplem, including the raw vibration waveform (Figure 3B, Supplementary Figure 4). AraVibS is an extended version of AraVib for the efficient detection of mutants using the *ωd* data, which was calculated by AraVib, and growth traits (Figure 1). In addition, the AraVib_model_control program that assists in creating a model for mutant detection by AraVibS is included (Figure 1). Figure 3A shows the structure of the expanded AraVib folder.

The calculated *ω_d_* is used for the analysis by itself (Figures 4A–C, 5A–B) and for the analysis combined with growth traits (Figure 5C–D, Supplementary Figures 5–7). Because the distribution of the combination of *ω_d_* and growth traits is different between wild type and tested cell-wall mutants (Figure 5C–D), we developed an extended version of AraVib for mutant detection and named it AraVibS (Figure 6A–C). To detect mutants, a multiple linear regression model and fitting to the normal distribution are used in this program (Figure 6A–B). To easily optimize the model under each condition, we also developed the AraVib_model_control program that assists in creating a learned model and in calculating parameters fitted to the normal distribution (Figure 6B–C).

**Figure 4.**
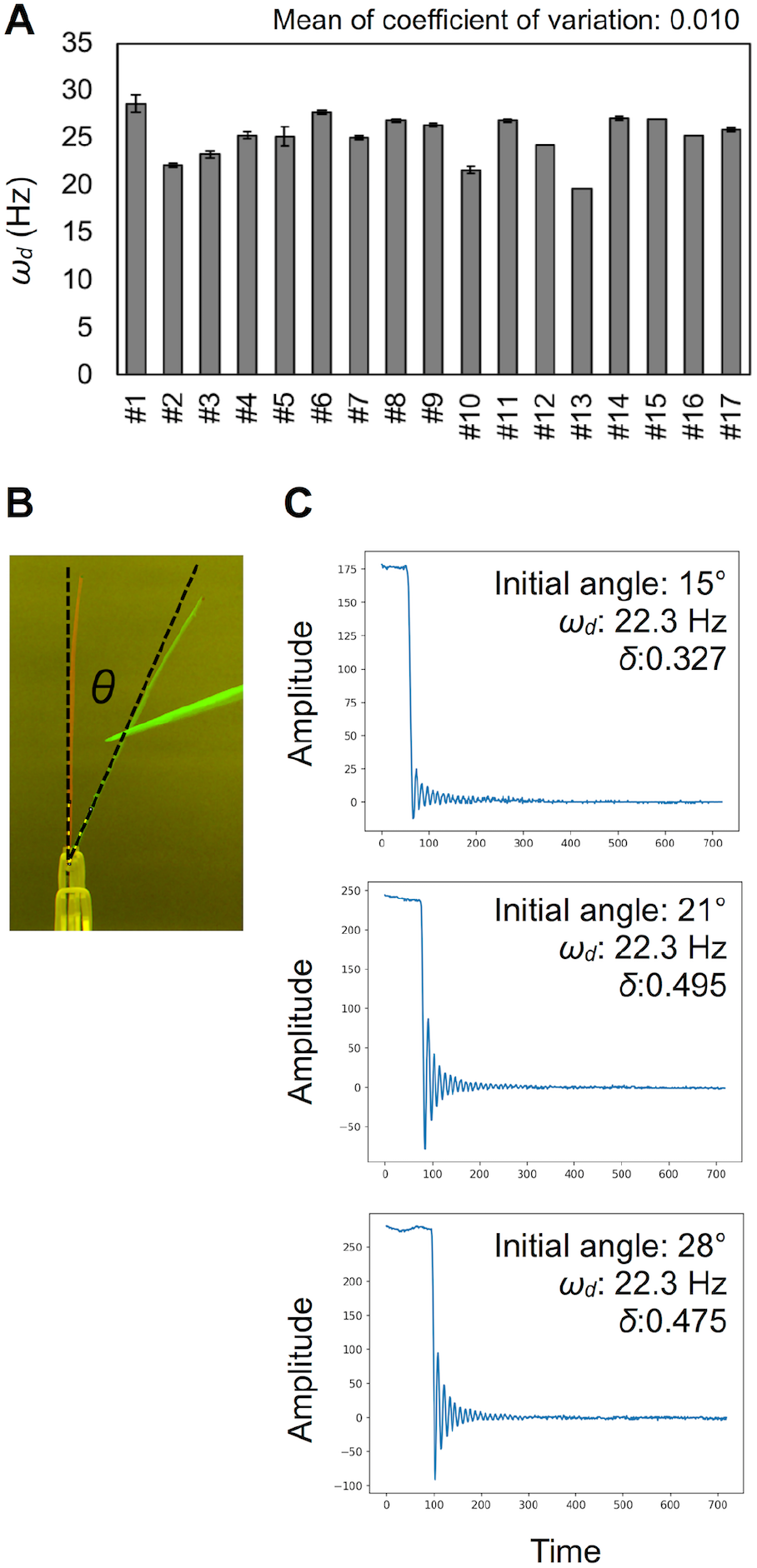
Validation for measurement accuracy of the damped frequencies *ωd*. (A) Deviations of *ω_d_* (Hz) in technical triplicates. Results of 17 independent wild-type plants (#1−#17). Data represents mean ± standard deviation. (B) The initial angle θ shown in a merged image. (C) Examples of vibration data showing the effect of the initial angle. See also Table 1.

**Figure 5.**
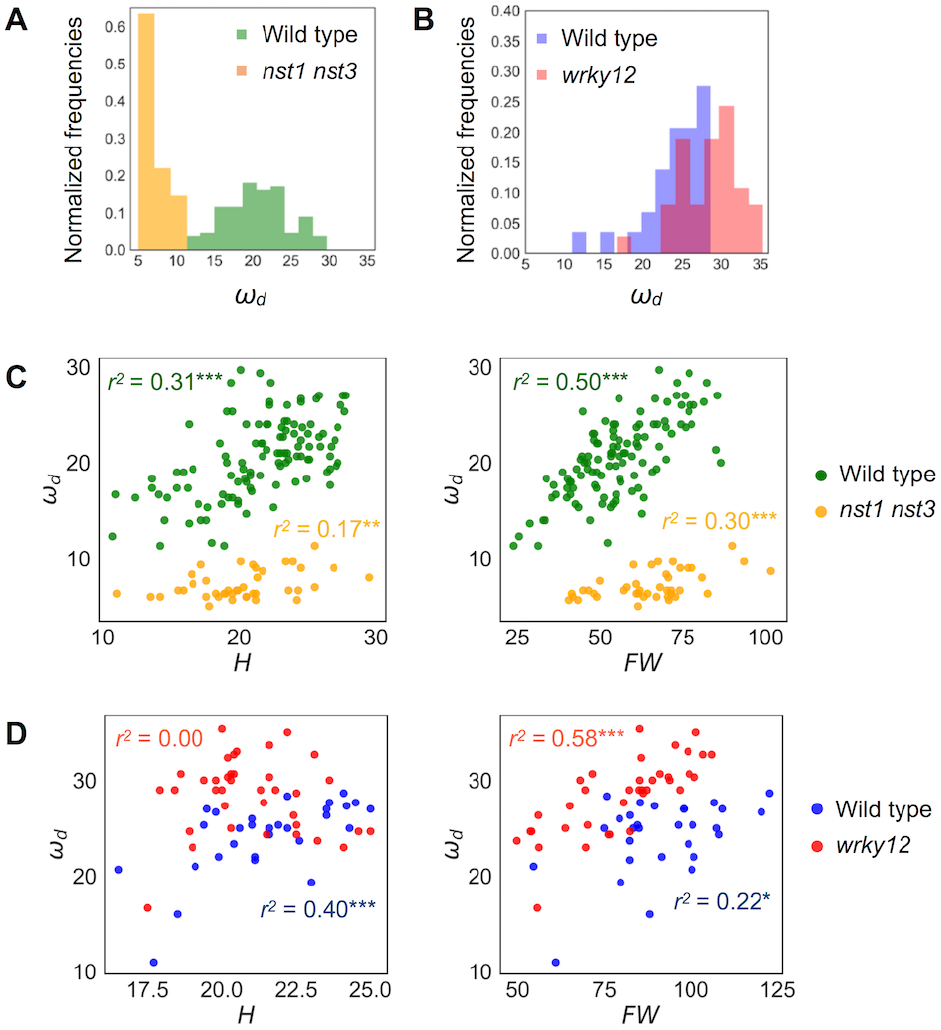
Distribution of *ωd* of wild type and cell-wall mutants. (A, B) Histogram of *ω_d_* (Hz). (C, D) 2-D scatter diagram between *ω_d_* (Hz) and H (cm) and FW (mg). *r^2^* means the square of Pearson’s correlation coefficient. \**p* < 0.05, \*\**p* < 0.01 and \*\*\**p* < 0.001. Samples of (A) and (C) were grown in condition 1 and of (B) and (D) were grown in condition 2. These samples correspond to those used to calculate the values shown in Table 3. n = 111 (wild type in A, C in green), 41 (*nst1 nst3* in A, C in orange), 29 (wild type in B, D in blue) and 37 (*wrky12* in B, D in red).

**Figure 6.**
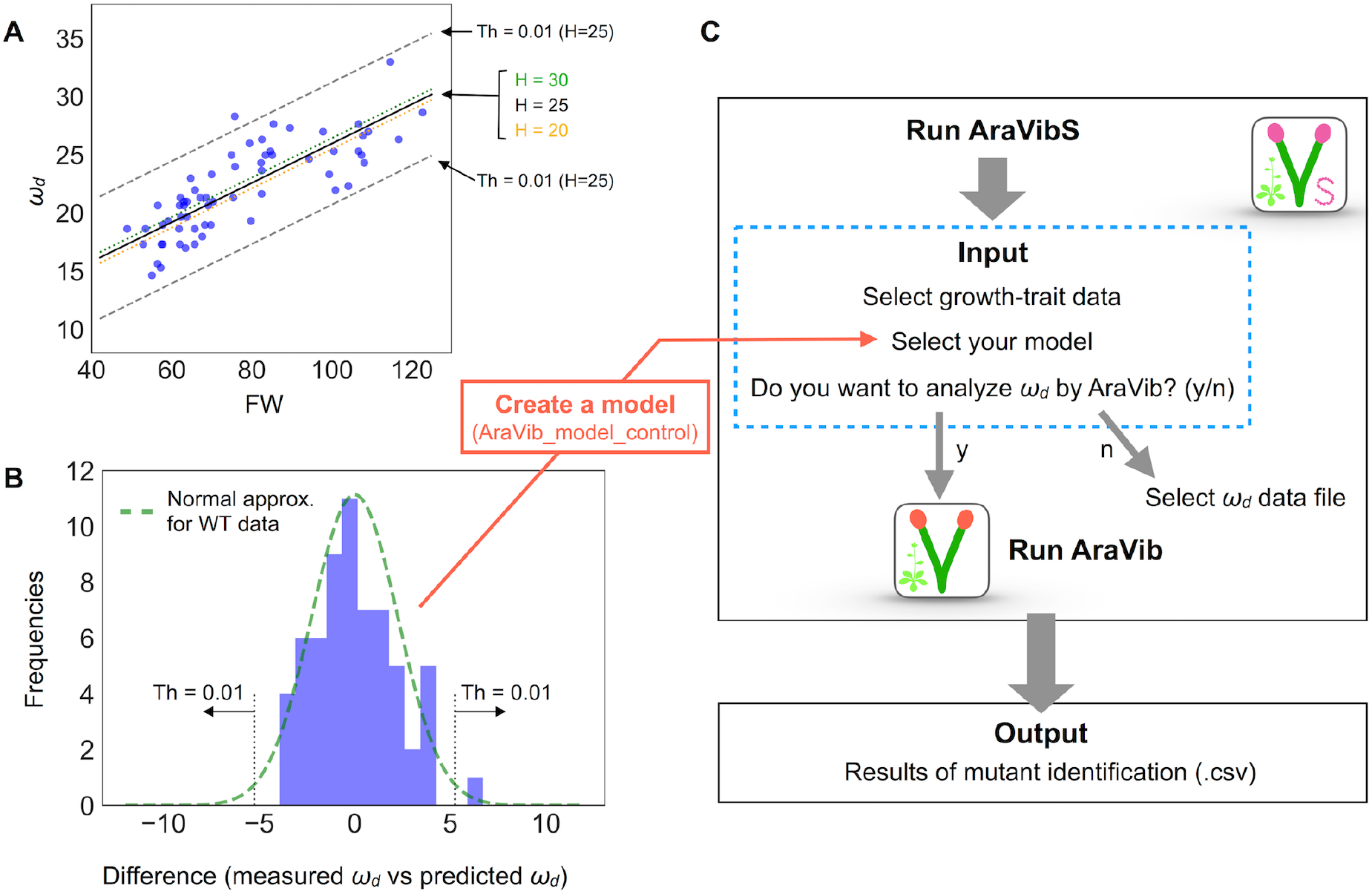
Model establishment by multiple linear regression and analysis diagram of AraVibS. (A) Regression line of *ω_d_* (Hz), FW (mg) and H (cm) of wild-type stems by multiple linear regression drawn on 2-D plot of *ω_d_* and FW (a black solid line, a green dotted line and an orange dotted line). The boundaries when the threshold value of *p*-value at H = 25 is 0.01 (shown as “Th = 0.01”) are indicated by gray dashed lines. (B) A histogram of the difference between the measured value and the predicted value of *ω_d_*. A green dotted line represents the fitted normal distribution. The boundaries when the threshold value of *p*-value is 0.01 (shown as “Th = 0.01”) are indicated by gray dotted lines. The number of samples is 63. (C) Analysis diagram of AraVibS. Whether to use AraVib or analyzed *ω_d_* file can be chosen. The output is the csv file of the result of mutant identification.

### Equations for estimating mechanical properties

By assuming that the stem is a simple spring-mass-damper system (Figure 7A), stiffness *k* and damping coefficient *c* can be estimated from *ωd* and logarithmic decrement (*δ*; Zebrowski, 1999; Villibor et al., 2016; Figure 7B). According to cZebrowski (1999) in tapered stems of Triticale and Triticum, the dynamic behaviors of the vertical columns were different from those of the horizontal beams. However, the difference was very small when only 20 cm of the basal portion was used or when the tip load was very light (Zebrowski, 1999). Therefore, we concluded that this model could be applied to the estimation of mechanical properties when the non-tapered part of the stem is used and the tip weight is close to zero.

**Figure 7.**
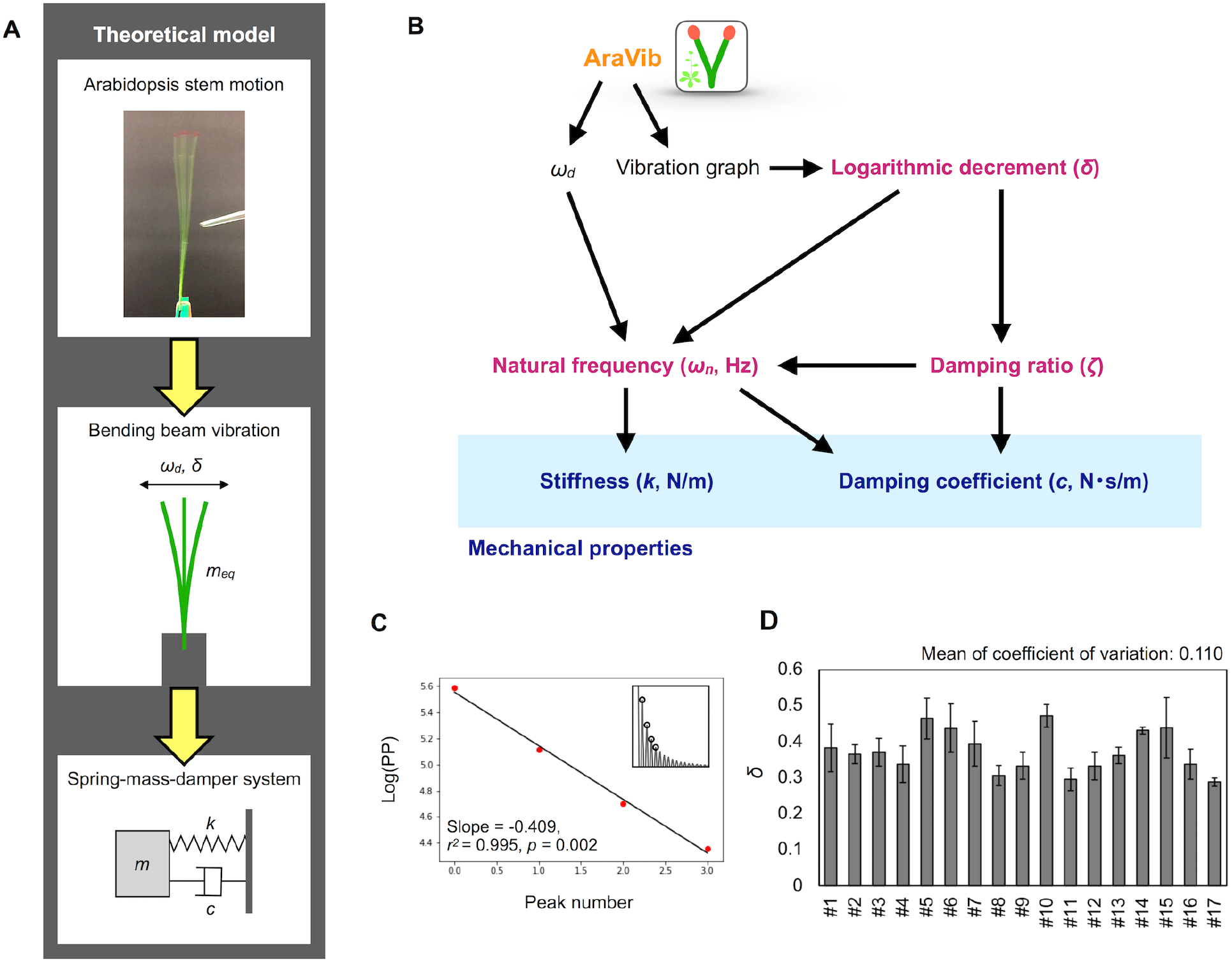
Mathematical model and calculation flow chart for estimation of mechanical properties. (A) Theoretical model of the motion of Arabidopsis stem. The motion is modeled as the spring-mass-damper system. According to this model, stiffness *k* and damping coefficient *c* are estimated. (B) Calculation flow chart for estimation of mechanical properties from the outputs of AraVib. (C) The graph for calculation of *δ*. “Log(PP)” represents the natural log transformed pixel value of the maximum peaks. Red points indicate the natural log transformed pixel values of the four consecutive maximum peaks used for calculation of *δ*, corresponding to the circles at the insets. Black lines indicate linear regression lines. (D) Deviations of *δ* in technical triplicates. Results of 17 independent wild-type plants (#1−#17). Data represents mean ± standard deviation.

The *δ* value was calculated from the maximum peak amplitude in each period in the graph of the raw vibration waveform (Zebrowski, 1999; de Silva, 2006; Figure 7C), as follows:

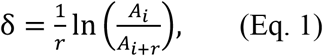

where *A_i_* and *A_i+r_* represent the maximum amplitudes of the *i* and *i+r* periods, respectively. The peak amplitude in each period was obtained by another program to automatically detect the maximum peaks from a graph of vibration data without Hanning-window processing. *δ* was estimated by a linear regression using multiple peak values to alleviate the influence of measurement errors. The pixel values of the peaks were natural log transformed and plotted for each peak number (Figure 7). A linear regression was performed on four possible consecutive data points from the first to ninth peaks, and the slope, *r^2^*, *p*-values were calculated. Among the calculated regression formulae, the absolute value of the slope of the regression equation where *r^2^* was maximum and the *p*-value was < 0.05 was adopted as *δ* (Supplementary Table 1). After calculation, the graphs of logarithmic plots of the pixel values of the peaks and the regression lines were visually confirmed. For samples with peak values that did not monotonically decrease or fit to the regression line, manual detection was performed using ImageJ (Abràmoff et al., 2004), and then, *δ* was calculated.

Based on two previous reports (Zebrowski, 1999; Villibor et al., 2016), the damping ratio (*ζ*), the natural frequency (*ω_n_*), the damping coefficient (*c*) and the stiffness (*k*) were estimated from *ω_d_* and *δ* (Figure 7B), as shown below.

*ζ* was estimated by the following equation.

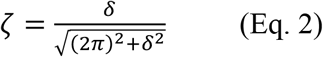

*ω_n_* was estimated by the following equation.

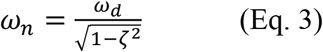

*c* was estimated by the following equation.

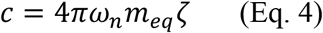

where *m_eq_* represents the equivalent mass of the oscillator (Figure 7). *m_eq_* is expressed by the following equation as described in Zebrowski (1999):

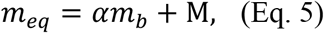

where M represents the mass of the attached tip-point weight, but in our system this is substantially zero (see the following section). Also, *m_b_* represents the weight of the beam; *α* = 0.245 in the case of a beam showing a uniform weight distribution. The basal stem portion (10 cm from the base) of *A. thaliana* inflorescences without any accessories showed a uniform weight distribution on average (Supplementary Figure 8). Of the 10-cm stems, 1 cm was used to fix to the setting board (see the following section). Thus, the remaining 9-cm section contributed to the vibration. Thus,

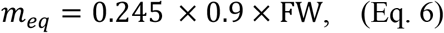

where FW represents the fresh weight of the 10-cm stem. The relationship between the natural frequency and the stiffness *k* was expressed by the following equation.

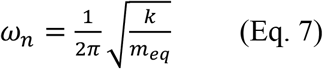

Thus, the stiffness *k* was estimated by the following equation.

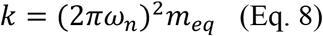

### Statistical analyses

For individual parameters, the statistical functions of the Pandas, Numpy and Scipy were used. Global scatter plots of all parameter pairs were drawn by the function of the Seaborn library. Histogram and scatter plots were drawn by the function of the pyplot package of the Matplotlib. For the statistical tests for differences, normality of data distribution (growth traits and oscillatory properties in each genotype and each growth condition) was tested using the Shapiro–Wilk Normality Test, and the homogeneity of variances was first analyzed with Bartlett’s test. Normally distributed data with equal variances between two groups were examined using two-tailed Student’s t-test. When the data was not normally distributed, a two-tailed Mann–Whitney U test was used. The methods of multiple comparison test are shown in each table (Tables 1, 2).

**Table 1.**
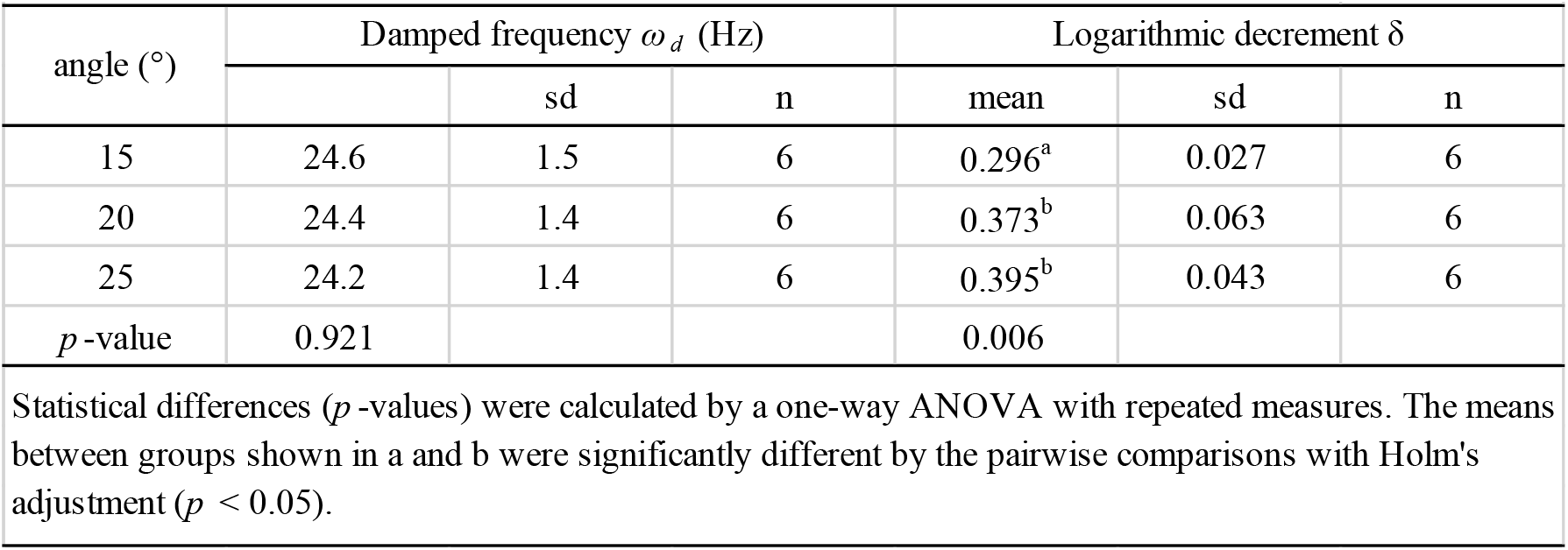
The effect of the initial angle on damped frequency and logarithmic decrement.

**Table 2.** The effect of the number of nodes on damped frequency and logarithmic decrement.

|  |  |  |  |  |  |  |
| --- | --- | --- | --- | --- | --- | --- |
| nodes | Damped frequency $\omega_d$ (Hz) | | | Logarithmic decrement $\delta$ | | |
|  | mean | sd | n | mean | sd | n |
| 1-2 | 24.2 | 3.1 | 7 | 0.354 | 0.095 | 6 |
| 3 | 24.8 | 2.4 | 11 | 0.397 | 0.060 | 11 |
| 4-5 | 24.8 | 2.4 | 7 | 0.340 | 0.040 | 6 |
| *p*-value | 0.976 |  |  | 0.233 |  |  |
| Statistical differences (*p*-values) were calculated by the Kruskal-Wallis H-test. |  |  |  |  |  |  |

### Preparation of alcohol-insoluble residues (AIRs)

To obtain AIRs, the red marker on the samples was wiped off after imaging of the stem vibration and the stem samples were soaked in 100% methanol for at least 1 d. The samples immersed in methanol were then treated sequentially with 100% acetone, methanol/chloroform (1:1) and 100% ethanol, and then dried overnight. AIRs of the samples were weighed with a precision balance.

### Other analyses

To calculate the initial angle, the images taken at the vibration start and after the vibration termination were merged. On the merged image, the angle was manually calculated by the function of ImageJ. For the histological analysis, we prepared hand-cut sections of the 1.5-cm base portion of the stem and observed the brightfield and UV autofluorescence using an Axioskop2 fluorescent microscope (Carl-Zeiss Inc.). The cross-sectional area of the stem was measured by ImageJ.

## Results

### Effect of the initial angle and the number of nodes on *ω_d_*

To calculate the *ω_d_*, we developed the experimental method and the AraVib software to analyze the free vibrations of stems fixed at their basal ends. First of all, we investigated the magnitude of the technical variations of *ω_d_* in the inflorescence stem of *A. thaliana* wild type. Technically triplicate experiments revealed that the variation of *ωd* was only ~1% (Figure 4A). Next, we examined how the extent of the initial stem angle (shown in Figure 4B) affects *ω_d_*. When the initial angle was roughly 20 to 30°, a typical waveform of damped oscillation was shown, while if the initial angle was ~15°, an abnormal waveform was observed (Figure 4C). Nevertheless, experiments in which each sample was tested at multiple initial angles indicated that *ωd* was nearly constant, regardless of the initial angle (Table 1). A previous study using the stem of *Equisetum hyemale* demonstrated the effect of nodes on stem vibration (Zajaczkowska et al., 2017). Therefore, we also examined the effects of the numbers of nodes on *ωd* in 10-cm stems of *A. thaliana*. We found that the number of nodes did not affect *ω_d_* (Table 2).

### Distribution of *ω_d_* in wild type, *nst1 nst3* and *wrky12*

Because the stem’s stiffness affects *ω_d_* in principle, we investigated whether the *ω_d_* values in two cell-wall mutants differed from that of the wild type. These mutants were the *nst1 nst3* mutant in which the mechanical strength was reported to be weakened (Mitsuda et al., 2007) and the *wrky12* mutant in which the mechanical strength was presumed to be altered because of ectopic formation of the secondary cell walls (Wang et al., 2010). Examples of the vibration waveforms are shown in Figure 3B and Supplementary Figure 4, and the results of the multiple-sample analysis are shown in Table 3. The growth traits of the measured samples are shown in Supplementary Table 2.

**Table 3.**
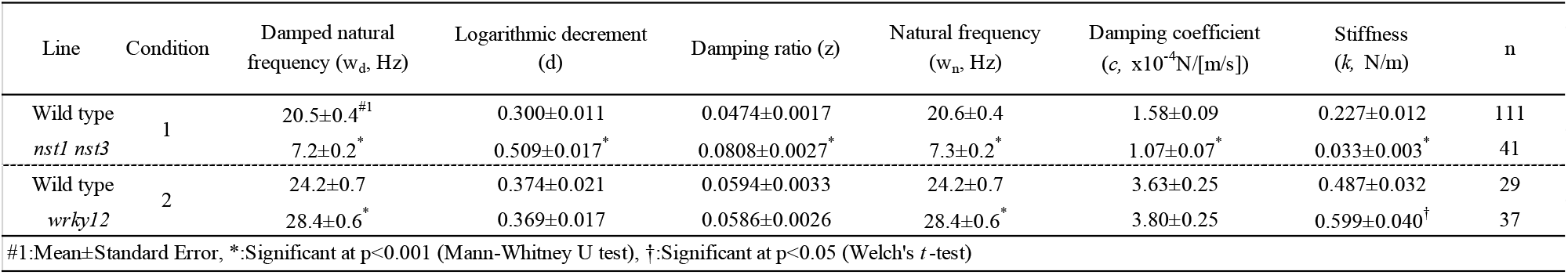
Oscillatory and mechanical properties of wild type and *wrky12* stems modeled as a vertically fixed and uniform prismatic beam without the oscillator.

The stem vibration waveform was a typical damped oscillation, regardless of genotype (Figure 3B, Supplementary Figure 4). However, the vibration of *nst1 nst3* was apparently slower (Supplementary Movies 1, 2, 5, 6), and the periods of the waves were larger than those of wild type (Figure 3B, Supplementary Figure 4). The major frequency, regarded as *ω_d_*, of the *nst1 nst3* stems was lower than that of wild-type stems according to the FFT analysis (Figure 3B, Supplementary Figure 4). The mean of *ωd* significantly decreased to 34%–36% of wild type in *nst1 nst3* (Table 3), and the distributions of the two genotypes were well separated (Figure 5). However, no significant differences in the speed of vibration or the forms of the damped oscillation curves between wild type and *wrky12* were found, although the FFT analysis indicated that the frequency of the *wrky12* stem was slightly higher than that of the wild type stem (Figure 3B, Supplementary Figure 4, Supplementary Movies 1, 2, 7, 8). The analysis of multiple samples revealed that *ω_d_* was significantly increased to 115%–120% of wild type in *wrky12* (Table 3), while the distributions of the two genotypes overlapped (Figure 5). This result suggests that *ω_d_* can be an indicator of changes in cell wall, which are accompanied by changes in stiffness.

### Identifying cell-wall mutants using the combination of *ω_d_* and growth traits

*ω_d_* showed a wide distribution in both wild type and *wrky12* mutants. It was assumed that the stiffness depended not only on the genotype but also on the degree of stem growth. To validate this assumption, we examined the correlation between *ω_d_* and two growth traits, H and FW, in wild type, *nst1 nst3* and *wrky12*. In a two-dimensional (2-D) plot, *ω_d_* showed a statistically significant positive correlation with H and FW under all conditions and in any genotype (Figure 5A, B). Thus, the stiffness also depends on the degree of the stem growth in *A. thaliana*.

The distribution of *wrky12* was well separated from that of the wild type on the 2-D plot of *ω_d_* and growth traits (Figure 5B). Therefore, we hypothesized that cell-wall mutants could be distinguished from the wild type group using the correlation between *ω_d_* and growth traits. Because *ω_d_* was dependent on the degree of stem growth, an equation to obtain the expected value of *ω_d_* from the H and FW values was determined by a multiple linear regression (Figure 6A). The FW value contributed more to *ω_d_* than the H value (Figure 6A). The 2-D plot suggested that the deviation from the regression equation of the wild type distribution fell within a certain range (Figure 6A). Based on the H and FW values, we calculated the expected value of *ω_d_* from the regression equation and estimated the deviation from the measured value. The distribution of the deviation could be fitted to the normal distribution (Figure 6B). Using the fitted result as a model, we developed AraVibS, which combines AraVib with a program that classifies whether the sample is a mutant by determining the difference from wild type using the *p*-value (Figure 6C). When the *p*-value threshold of the model was set to 0.01 in each experimental condition, 100% of *nst1 nst3* samples were specified as mutant, while 60.0% of *wrky12* samples were specified as mutant (Table 4).

**Table 4.**
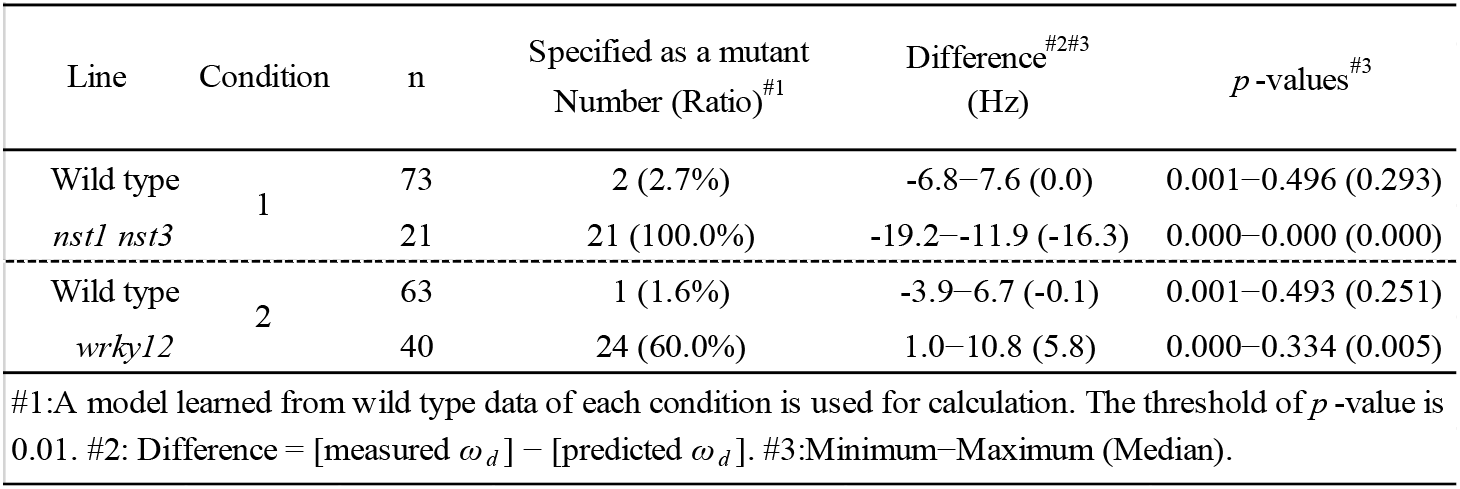
The result of mutant detection by AraVibS.

### Quick estimation of the mechanical properties from the AraVib output

Using the AraVib output, we attempted to calculate *k* and *c*. As mentioned in the Materials and Methods, *k* and *c* are calculated from *ω_d_* and *δ* (Figure 7A, B). *δ* can be estimated from the vibration waveform (Figure 7C). As is for *ω_d_*, technical variations of *δ* were investigated. The variation of *δ* was more than 10% in technical triplicates (Figure 7D). *δ* was consistent when the initial angle is in the range of 20 to 30° but tended to decrease when the initial angle was ~15° (Table 1, Figure 4C). The number of nodes had limited effects on *δ* (Table 2).

The estimated mechanical properties (*k* and *c*) are shown in Table 3, along with *ω_n_* and *ζ*. The difference between *ω_n_* and *ω_d_* was approximately 1/10 Hz in both tested genotypes, which was below the frequency resolution (1/3 Hz) of *ω_d_*. Thus, the *ω_n_* and *ω_d_* values were almost the same (Table 3). The *k* value was significantly reduced to 13%–16% of wild type in *nst1 nst3* (Table 3), and the distribution was separated (Figure 8). The *c* value was significantly lower in *nst1 nst3* (63%–72% of wild type), but the distribution overlapped with that of wild type (Figure 8A). While the mean of *c* in *wrky12* was almost equivalent to that of wild type, the *k* value significantly increased to 115%–131% of wild type (Table 3). The distribution of *k* overlapped between wild type and *wrky12* (Figure 8A).

**Figure 8.**
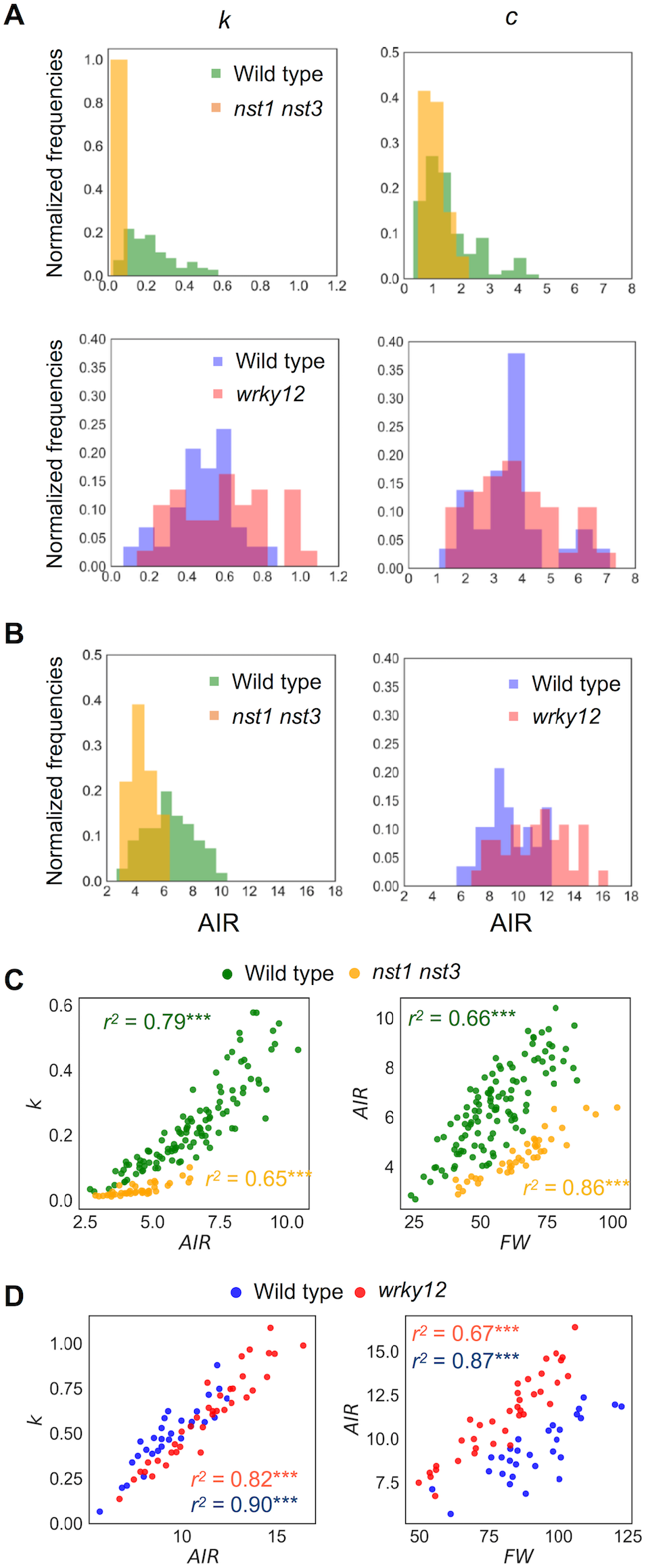
Distribution of mechanical properties and AIR. (A) Histogram of mechanical properties, *k* (N/m) and *c* (×10-4Nm-1s-1). (B) Histogram of AIR (mg). (C, D) 2-D scatter diagram of *k* (N/m) and AIR (mg) or FW (mg) and AIR (mg). *r^2^* means the square of Pearson’s correlation coefficient. \*\*\**p* < 0.001. These samples correspond to those used to calculate the values shown in Table 3. n = 111 (wild type shown in green), 41 (*nst1 nst3* shown in orange), 29 (wild type shown in blue) and 37 (*wrky12* shown in red).

### The cell-wall density in the 10-cm stem was well correlated with *k*

The *k* value drastically decreased in *nst1 nst3* and slightly increased in *wrky12* compared with wild type. In *nst1 nst3*, the amount of cell wall decreased because the secondary cell wall is absent (Mitsuda et al., 2007; Sakamoto and Mitsuda, 2015), whereas in *wrky12*, ectopic wood formation occurs in the pith (Wang et al., 2010) and the amount of cell wall is assumed to increase. Measurements revealed that the cell-wall density (AIR/FW) significantly decreased to 59%–60% of wild type in *nst1 nst3* and significantly increased to 131%–137% of wild type in *wrky12* under our experimental conditions (Figure 8B, Supplementary Table 2).

In the 2-D plot, a clear positive correlation between *k* and AIR was observed (Figure 8C, D). A regression analysis revealed the strong positive correlation between *k* and AIR (*r^2^*: wild type, 0.79 or 0.82; *nst1 nst3*, 0.65; *wrky12*, 0.90), which was comparable with the correlation between AIR and FW (*r^2^*: wild type, 0.66 or 0.67; *nst1 nst3*, 0.86; *wrky12*, 0.87). These results support a close relationship between the cell wall density and *k*.

### Comparison of the *ω_d_* and mechanical properties of wild type populations grown under different conditions

There were differences in the distributions of *ω_d_* and mechanical properties between wild-type samples grown under two conditions (Table 3, Figure 9A, B). In addition, the distributions of FW and AIR changed dramatically (Figure 9A, B, Supplementary Table 2). While wild type samples grown on the two conditions showed overlapping distributions on the 2-D plots of *k* and AIR, the distributions appeared different on the 2-D plots of *ω_d_* and FW. To investigate the differences associated with these two growth conditions in more detail, we observed stem cross-sections.

**Figure 9.**
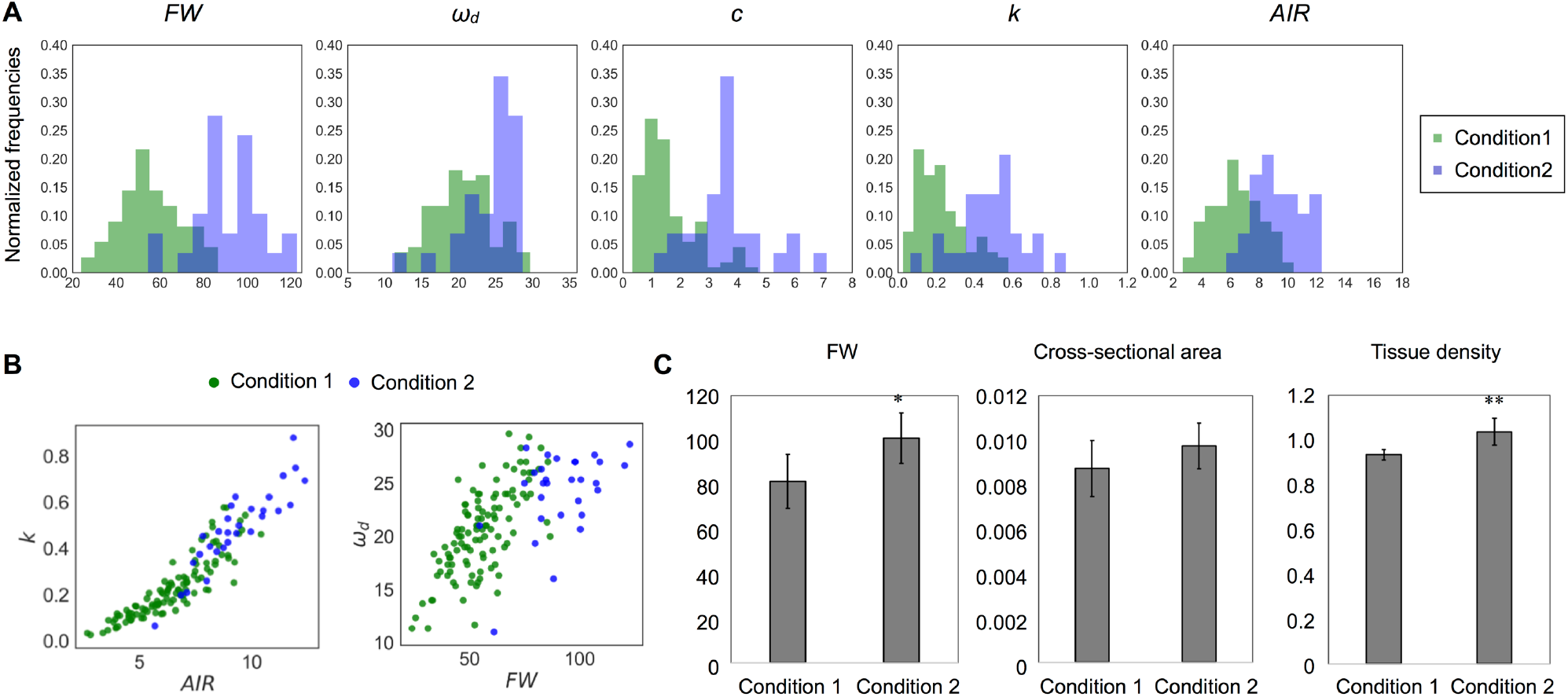
Difference of distribution of each parameter between different growth conditions. (A) Histogram and (B) 2-D plots of AIR and *k* or FW and *ω_d_*. All parameters appear to be sensitive to change depending on growth conditions. (C) The 10-cm FW (mg), cross-sectional area (cm2) and the tissue density (FW/Volume, ×10-2mg/m3) estimated from them. Data represent mean ± standard deviation. n = 5 (condition 1), 7 (condition 2). Sampling was performed for stems with a height of 20-25 cm.

There was no obvious histological difference between the stems of plants grown under these two conditions (Supplementary Figure 9) nor were the cross-sectional stem areas significantly different (Figure 9C). The tissue densities estimated using FWs and the cross-sectional areas were significantly different between plants grown under the two conditions (Figure 9C). Thus, the FW, AIR and tissue density of wild-type stems, as well as the *ω_d_* and *k* values, differed between plants grown under the two conditions.

## Discussion

In this study, we developed a high-throughput analysis method for the free vibrations of Arabidopsis inflorescence stems that can identify cell-wall mutants with altered stiffness and quickly estimate mechanical properties. The AraVib software for free vibration analysis outputs a *ω_d_* list for each analysis and three kinds of graphs for each sample. By analyzing *ωd* and growth traits with AraVibS, 100% of *nst1 nst3* and 60% of *wrky12* samples were correctly judged as mutants. Furthermore, mechanical characteristics were estimated using *ω_d_* and the graph of the vibration waveform.

Experiments with free vibration generally take a short time to capture vibrations that decay quickly. Because the time for data acquisition affects the analysis throughput, it is a fundamental advantage of the free-vibration method. However, this method uses one fixed end (termed the cantilever beam), unlike the free-free vibration method, and thus may not reflect the *k* level of the whole stem because various factors, including clamping effect, influence free vibrations. However, we concluded that our method is sufficient to estimate mechanical properties quickly for the investigation of genetic factors, because it was able to detect changes in the stem’s stiffness of Arabidopsis mutants.

Another problem of measuring free vibration is that a high-speed camera or detector is required to obtain a sufficient resolution from the short-time displacement. Other studies of free vibration of plant stems used cameras with time resolutions of 200–1,000 fps (Der Loughian et al., 2014; Villibor et al., 2016; Zajaczkowska et al., 2017). In this study, we used high-speed shooting at 240 fps with the default function of an iPhone/iPad. According to the Nyquist–Shannon sampling theorem, for a frequency analysis without aliasing, the data sampling frequency must be more than twice the frequency of the measurement target (Nyquist, 1928; Shannon, 1949). Because the maximum *ωd* of the 10-cm inflorescence stem of *A. thaliana* was 35 Hz, and the frame rate was more than 6.8 times the frequency, it was sufficient for the analysis.

The most important factor for the throughput of our method is the AraVib software. AraVib can analyze many videos in batches with only a few operations in the command line. Because Python knowledge is not necessary, researchers who are not familiar with computer programs and image analysis can easily use the software. The throughput rate is 30–40 s per sample, and all movies shot in a day are automatically analyzed by the batch process, which is essential for a large-scale screening. In vibrational analyses based on videos, coordinates are obtained by tracking a specified target. In previous studies, to capture stem vibrations, the features were tracked by an automatic analysis using the Kineplant-CR toolkit in MATLAB (Der Loughian et al., 2014), by a semi-automatic analysis with a free software for tracking (Zajaczkowska et al., 2017) or by a manual analysis (Yazdanmanesh and Kalantari, 2015). In Spatz and Speck (2012), vibrations were captured by detecting tape, as a small marker, with the commercial software. Although it is not certain whether the image analysis was batch processed or *ω_d_* was calculated seamlessly in these studies, the numbers of tested samples were relatively small and they did not appear to have high-throughput potential (Spatz and Speck 2012; Der Loughian et al., 2014; Yazdanmanesh and Kalantari, 2015; Zajaczkowska et al., 2017). Currently, the highest throughput method for determining free vibration uses a custom-made device rather than a camera (Żebrowski, 1991; Żebrowski, 1999). Because this custom-made device includes a photodiode, amplifier and time-counter, it is not considered widely attainable for many biologists. In this study, more than 300 samples were analyzed, which is much more than in other studies using a camera. In addition, more than 50 samples could be analyzed in under 2 h. Therefore, this is the most high-throughput method compared with others available to many biologists.

We also developed AraVibS, which automatically detects mutants. As mentioned above, AraVibS was able to detect 100% of *nst1 nst3* and 60% of *wrky12* as mutants. AraVibS uses a multiple regression equation that calculates the expected value of *ω_d_* from H and FW as a learned model, but it should be used carefully. Based on our analysis, the distributions of FW and *ω_d_* changed even in the wild type population depending on growth conditions. This difference is likely caused by the differences in FW, AIR and tissue density. In fact, tissue density affects tissue strength (Niklas, 1993), which may result in changes in *ω_d_*. When conducting mutant detection using AraVibS under new conditions, data from the wild type or control line should be analyzed under the same experimental conditions with AraVib_model_control.

Using *ω_d_* and *δ* calculated from the output of AraVib, *k* and *c* were estimated according to the spring-mass-damper system model. The technical variation of *ω_d_* was approximately 1% and was stable, independent of both the initial angle and the number of nodes. However, the variation of *δ* was approximately 11% in technical triplicates and was affected by the initial angle. For the calculation of *k*, the contribution of *δ* is very small, resulting in the estimated error of *k* being approximately 2%. For the calculation of *c*, the contribution of *δ* is not ignorable, which results in an estimated error of *c* that is much larger than that of *k*. In *nst1 nst3*, based on the *k* value, stiffness was greatly decreased to approximately 15% of the wild type, which was consistent with our previous results in which the elastic modulus of *nst1 nst3* in the longitudinal direction was greatly reduced (Mitsuda et al., 2007; Yoshida et al., 2013). Thus, our method appears reliable enough to compare the *k* values of the stems of wild type and *nst1 nst3*. In addition, there was a difference in *k* between wild type and *wrky12*. Although the mechanical properties of *A. thaliana wrky12* stems have not been reported previously, there may be a difference in *k* between wild type and *wrky12*. Additionally, there was a difference in *ω_d_* between wild type and *wrky12*. Considering that *ω_n_*, which is almost the same value as *ωd*, is inversely proportional to the square root of *m_eq_* (Eq. 6) and that *k* is strongly correlated with AIR, the vibrational analysis might be sensitive to changes in AIR/FW. However, more investigations will be needed to clarify this point.

A regression analysis revealed a strong and positive correlation between *k* and AIR. This strongly suggests that *k* may be closely related to the amount of components of secondary cell walls, which accounts for the majority of AIR. According to an analysis of wood, in the transverse direction, hemicellulose and lignin are inferred to contribute to the elastic modulus (Salmén, 2004). In addition, the relationship between secondary cell wall components and stem’s mechanical strength has been intensively studied using *A. thaliana*, poplar and tobacco. For example, a decrease in the total amount of lignin causes a decrease in *k* and increases the viscoelastic decay (Köhler and Spatz, 2002; Hepworth et al., 1998; Hepworth and Vincent, 1999; Köhler and Telewski, 2006; Özparpucu et al., 2017). Increasing the syringyl/guaiacyl ratio of lignin increases the modulus but does not change the bending stiffness (Köhler and Telewski, 2006). A decrease in xylan causes a decrease in wall thickness and causes a decrease in elastic modulus despite an increase in the relative amount of lignin (Li et al., 2011). However, an increase in the expression level of xyloglucanase increases the cellulose content and specific gravity, causing an increase in the elastic modulus (Park et al., 2004). Despite such intensive research, details regarding the relationship between cell wall components and mechanical properties remain unknown. A high-throughput method like ours enables the comprehensive investigation many genetic variations.

We demonstrated that our method is useful for the automatic detection of mutants and a quick estimation of mechanical properties. Among the existing screening methods used to evaluate the stem strength, visual observations can only detect drastic changes in strength, and the reliability of the hand-bending test strongly depends on the researcher’s skill. Although the application of this method is not as rapid as visual observations or the hand-bending test, this is a robust, quantitative and relatively sensitive method that does not require specific technical skills. The throughput rate of our method is comparable with or greater than that of screening by anatomical analysis with hand sectioning. Thus, we believe our method with further improvements will be suitable for large-scale screening studies.

## Author Contributions

MN contributed to conception of the study, implemented AraVib and AraVibS, performed data collection and analysis, and wrote the initial draft of the manuscript. MN and MT contributed to the design of AraVib and AraVibS. MN, MT and KY contributed to interpretation of data. MN, MT, SS and NM contributed to the design of the laboratory experiments. MN, MT, KY and NM contributed to manuscript revision. All authors read and approved the submitted version.

## Abbreviations

*ω_d_*: the damped natural frequency
*δ*: the logarithmic decrement
*ζ*: the damping ratio
*ω_n_*: the natural frequency
*c*: the damping coefficient
*k*: the stiffness
*m_eq_*: the equivalent mass of the oscillator
FW: the fresh weight of the trimmed stem
AIR: the alcohol-insoluble residues of the trimmed stem
H: stem height
*FFT*: Fast Fourier Transform

## Acknowledgments

We thank A. Hosaka, M. Yamada and Y. Sugimoto for technical support. This study was performed under the support of JST ALCA program (to N. Mitsuda). We thank Lesley Benyon, PhD, from Edanz Group (www.edanzediting.com/ac) for editing a draft of this manuscript.

## Availability

The GitHub page for the AraVib project is https://github.com/MTNakata/AraVib.

The free Python packages or libraries, which are used in this study, are the OpenCV-Python package (https://pypi.python.org/pypi/opencv-python), NumPy package (https://pypi.python.org/pypi/numpy), the Python Pandas package (https://pypi.python.org/pypi/pandas/0.18.1/), SciPy library (https://pypi.python.org/pypi/scipy), the pyplot package of the Matplotlib (https://pypi.python.org/pypi/matplotlib), Scikit-learn package (http://scikit-learn.org/stable/index.html) and the Seaborn library (https://pypi.python.org/pypi/seaborn).

